# A classification of functional pitcher types in *Nepenthes* (Nepenthaceae)

**DOI:** 10.1101/852137

**Authors:** Martin Cheek, Matthew Jebb, Bruce Murphy

## Abstract

We classify *Nepenthes* species into 12 functional pitcher types, based on combinations of traits that appear to comprise different syndromes for capturing nutrients, usually from animals. For nine of these types the trapping syndromes are already documented, six targeting live animals (hence carnivorous), and three targeting other nutrient sources (non-carnivorous). Yet, for three pitcher types here is no previous documentation of the syndrome and we do not yet know what sources of nutrients are being targeted. Mapping all these pitcher types on the latest, near comprehensive species-level phylogenomic tree of *Nepenthes* (Murphy et al. 2019) shows that apart from the ancestral pitcher type 1, most of the remaining pitcher types have evolved independently, in different parts of the phylogenetic tree, usually in several different places. Each of the 12 pitcher types is characterised morphologically and illustrated, its trapping syndrome discussed, and example species are given. An identification key to the 12 pitcher types is presented. The possibility of additional pitcher types being present is discussed.

## Introduction

The pitchers of Nepenthes come in many shapes, sizes, and colours. Within a single individual these features vary, sometimes dramatically, as the plant matures. One of the most characteristic features of these plants is the dimorphy of pitchers in most species. These are distinguished as upper and lower pitchers, occasionally with distinct intermediate forms. The switch from one form to another is associated with the onset of elongate stems and flowering (Jebb, 1991). It is the upper pitchers (or in those few species that do not form upper pitchers, the intermediate or lower pitchers) that usually characterise a species, possessing the features that enable it to be identified and separated from other species.

Historically the presumption of insectivory meant that little attention was paid to the subtlety of pitcher shape and form in this regard. It is now recognised that different species of *Nepenthes* can obtain their nutrients from several different, mainly animal, sources, using a diversity of mechanisms and features or characters present in their pitchers, that therefore appear to be “functional traits” (i.e. Bauer et al. 2012a).

Based on such traits the pitchers of the known species of *Nepenthes* are classified here into 12 different functional types. Nine of these types correspond to different, more-or-less well-documented mechanisms for obtaining nutrients. But, for three of these types we can only conjecture what mechanisms might be used and what sources of nutrients might be targeted.

Two of the 12 types have only a single documented species, such as type 10, represented only by *N. albomarginata* Lobb ex Lindl. At present this is the only species known to specialise exclusively in trapping termites, which it entices by means of the properties of the white hairy band on the outside of the pitcher below the peristome. No other species has such a band. However, ten of the 12 types have two to many species, and with the exception of the “ancestral” pitcher type 1, they are scattered as individuals or in clusters, through the evolutionary tree of Nepenthes (Murphy et al. 2019). It is therefore clear that nine of the 12 types documented here have each arisen independently more than once.

## Materials and Methods

Following a review of the literature on *Nepenthes* trapping syndromes, the species of the genus were classified into nine discrete morphological groups (each linked to a different syndrome) based on the functional traits present in their latest stage, usually upper, pitchers. Those species not falling into these nine groups fell into three further distinct morphological groups (pitcher types 4, 6 and 12) which were also characterised. An identification key to the 12 pitcher types was constructed using conventional taxonomic methods. Morphological data (e.g. pitcher shape and proportions) were taken from Jebb & Cheek (1997) and Cheek and Jebb (2001) and for species published subsequently, the protologues (e.g. Cheek & Jebb 2013a-h). Presence or absence of a “conductive” or waxy zone in the pitchers, not usually recorded in taxonomic literature, was obtained by study of reference herbarium specimens at the Kew Herbarium (K), and in some cases where specimens were not available, by reference to photographs available in McPherson (2009). Waxy zones have an opaque, dull, off-white appearance, while non-waxy surfaces have a glossy appearance. Presence or absence of visco-elastic versus watery pitcher fluid, also not usually recorded in taxonomic literature, was obtained from observation of late stage pitchers from live plants of 30 representative species in the Tropical Nursery of the Royal Botanic Gardens, Kew, U.K. in Nov. 2019, and (one species) at the National Botanic Gardens, Ireland (Table 1), also from the literature cited in the accounts of pitcher types 1-12 below. Visco-elastic fluid was detected in the species listed in table 1, by inserting two fingers into the pitcher fluid, withdrawing them from the fluid and then separating the two fingers. In visco-elastic species a filament is developed between the fingers, but is absent in non-visco-elastic species. The figures representing the 12 pitcher types are all drawn by Matthew Jebb from photos of live plants identified using Cheek & Jebb (2001).

## Results

A fundamental division between pitcher types is seen on the inner surface of the pitcher, between “waxy” (types 1, 7, 8, (9), 10) and “non-waxy” (types 2-6, (9), 11 & 12) species. It was Macfarlane (1908: 20) who first wrote in detail about waxy versus non-waxy zones and about the variation from one species to another. Bauer et al. (2012a) later documented this fundamental division in more detail (see below). The fundamental type of *Nepenthes* pitchers, type 1 of this classification, is seen in the earliest diverging, ancestral species such as *Nepenthes pervillei* Bl. and *Nepenthes distillatoria* L.f., but type 1 pitchers also predominate in Sect. Montanae Danser of the mountains of Sumatra and the Malay Peninsula e.g. *N. sanguinea* Lindl., and Sect. Pyrophytae Cheek & Jebb of Indo-china, e.g *N. smilesii* Hemsl. Most species of Sect. Alatae Cheek & Jebb (in the wide sense), from Luzon to Mindanao, also have type 1 pitchers.

### Key to functional pitcher types of Nepenthes (based on the latest stage pitchers developed)

1. Pitchers with a pale, matt, waxy (conductive) zone on the upper inner surface below the peristome, separated the from the glossy (detentive) zone at the base of the pitcher by an external visible raised line, the ‘hip’; liquid usually non-viscous ………………………………2 Pitchers entirely glossy on the inner surface, lacking matt, waxy zone, or with only a vestigial triangular remnant below the insertion of the lid, hip absent; liquid often (types 2, 3, 12 and *N. aristolochioides*) viscous …………………………………………………………………..6
2. Pitchers with a white hair band on the outer surface below the peristome ………………………………………………………………….. **Type 10, Termite Trap** Pitchers lacking a white hair band …………………………………………………………………..3
3. Lower surface of pitcher lid with either wax scales or hairs in the central portion ………………………………………………………………….. **Type 7. Flick of the lid** Lower surface of the lid lacking wax scales or hairs in the central portion………………………………4
4. Pitchers highly elongated, length: breadth ratio c. 10: 1 frequently acting as a day roost for bats ………………………………………………………………….. **Type 8. Bat-roost** Pitchers not highly elongated, length: breadth ratio <8: 1 not (or infrequently) acting as a roost for bats ………………………………………………………………….. 5
5. Pitchers with mouth lateral, vertical; apical windowed dome ………………………………**Type 9. Light-trap** Pitchers with mouth apical, horizontal, lacking an apical dome ………………………………**Type 1. Generalist**
6. Pitchers narrowly funnel-shaped …………………………………………………………………..**Type 2. Narrow-funnel** Pitchers ellipsoid, broadly cylindrical, domed, or broadly funnel-shaped ………………………………7
7. Pitchers often reclining, ellipsoid, usually massive and subwoody, diameter of mouth c. 10 cm; lid held erect of reflexed …………………………………………………………………..**Type 5. Tree shrew lavatory** Pitchers erect not reclining, not ellipsoid, but broadly cylindrical, domed or broadly funnel-shaped ………………………………………………………………….. 8
8. Pitchers broadly funnel-shaped, length: breadth ratio 1 to 1.5:1; usually 3-5 cm long, the upper part cup or bowl-shaped, wider than long, the lower part narrowly cylindrical ………………………………9 Pitchers broadly cylindrical or domed …………………………………………………………………..10
9. Pitchers lacking a very broad (to 3.5 cm wide), peristome absent or <1 cm wide …………………………………………………………………………………………… ………………**Type 3. Flypaper** Pitchers with a very broad (to 3.5 cm wide), flat peristome ………………………………**Type 12. Flat Lip**
10. Pitchers with mouth lateral, vertical, apical windowed dome ………………………………**Type 9. Light-trap** Pitchers with mouth apical, horizontal or inclined, lacking a windowed dome ……………………………… 11A
11. Lid held on a well-developed vertical column; outer surface of pitcher often with long, early caducous hairs …………………………………………………………………..**Type 6. Globose-hairy** Lid lacking a distinct column; peristome ridges not wing-like, hairs if present stellate ………………………………12
12. Pitchers never in carpets, with lid length: breadth 1.5-2: 1, held over the mouth, with nectar glands on the lower surface …………………………………………………………………..**Type 4. Stout Cylinder** Pitchers usually in carpets on the ground, with lid narrowly oblong length: breadth ratio c. 4:1, reflexed, not held over mouth, lacking nectar glands ………………………………**Type 11. Pitfall**

**Type 1 (Fig. 1).**
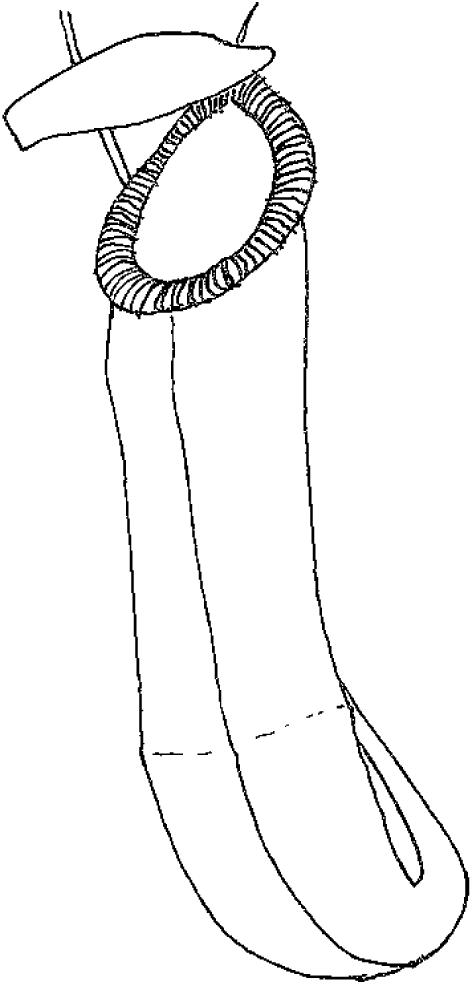
Ancestral (*N. mirabilis*) These species have upper pitchers which are more-or-less ovoid-cylindric in shape. The basal, ovoid part, often but not always a little wider than the upper part, is usually separated from the upper, usually cylindrical part by a “hip” or raised ridge that girdles the circumference of the pitcher on the exterior (also present in the six species of pitcher types 7, 8 and 10). Examination of the inner surface shows that the “hip” divides the glossy, broad, basal, fluid-containing, digestive, part (referred to as the “detentive” zone by Macfarlane (1908) because prey are detained here) from the upper opaque, dull, off-white to slightly purple, waxy or “pruinose” zone. This waxy zone is referred to as the “conductive” zone by Macfarlane because it helps conducts the insects downwards to the detentive zone with the digestive fluid. Such type 1 pitchers are shown in Fig. 1, exemplified by the upper pitchers of *Nepenthes mirabilis* (Lour.) Druce. The waxy zone is so-called because its surface is made up of numerous minute platelets of wax, each platelet like a roof-tile and held on a fragile stalk from the underlying surface. Juniper et al. (1989) documented how the waxy zone functions to prevent animals, usually insects, from leaving the pitchers once they have fallen in. Pressure from a foot placed on a wax “tile” will result in the stalk breaking, so that the insect loses its foothold and may fall back into the pitcher.

Type 1 species seem to derive their nutrition, so far as is known, by trapping a wide range of insects, as documented in detail by Jebb (1991) for *Nepenthes mirabilis* in Papua New Guinea. However, the number of prey individuals is dominated by ants: >80 % of prey in both upper and lower pitchers. More recent analysis of trapping syndromes in six species of *Nepenthes* in Brunei concluded that cylindrical pitchers with waxy walls (type 1) are particularly efficient at trapping and retaining ants, compared with other syndromes (Gaume et al. 2016). Earlier, Bonhomme et al. (2011) concluded that “wax only appears to be efficient for ants”.

**Type 2 (Fig. 2).**
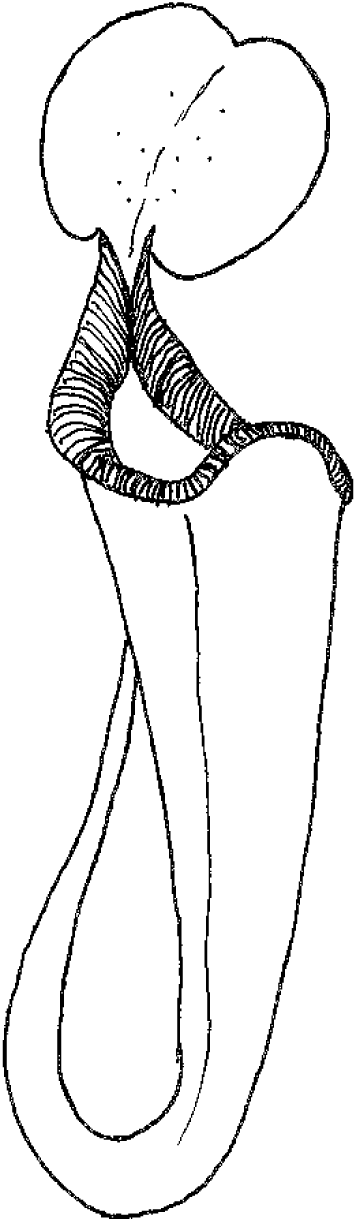
Narrow-funnel (*N. rafflesiana* Jack). In the upper pitchers of these and pitcher types 3-6 and 11-13 and in one species of type 9, the glossy detentive zone extends from the base of the pitcher to the peristome and no waxy layer can be seen, or just a vestigial remnant remains, such as a small triangle below the insertion of the lid. The hip is lost or is present as a remnant immediately below part of the peristome. The pitchers are usually narrowly funnel-shaped in type 2, with a length: breadth ratio of about 3 or 4: 1. This type can be seen in the upper pitchers of *N. rafflesiana, N. fusca* Danser, *N. maxima* Nees, *N. sumatrana* (Miq.)Beck, *N. copelandii* Merr. ex Macfarl., *N. treubiana* Warb. What is happening here, and why? Ulrike Bauer of the University of Bristol and co-authors (Bauer et al. 2012a) pointed out that the species which lack a waxy zone tend to have wider peristomes. Earlier they had shown how the peristome functions: becoming super-slippery only when wet (Bauer et al. 2008). Bauer et al. (2012a) inferred that these non-waxy species depend on the added efficiency of their wider-than-usual peristomes to send their prey into the inside of the pitcher. Bauer et al. conjectured that waxy species are less dependent on the efficiency of their generally narrower peristomes to send prey into their pitchers. Once prey enter, the waxy zone means that they cannot climb out. Using the molecular phylogenetic data of Meimberg et al. (2001), Bauer et al. (2012a) found that waxy zones are present in the most ancestral species and that non-waxy species have been derived from them multiple times. Moran et al. (2012) built further on the work of Bauer et al. (2012a). They considered a third trait or factor, beyond waxy/non-waxy and peristome broad/narrow. This is that several species of Nepenthes have “visco-elastic” (gloopy) pitcher fluid rather than watery fluid. Here, if the pitcher contents are poured out, they form a single elastic thread, as though containing glycerine. This feature is correlated to some extent with a number, but not all, non-waxy species.

Notably, we have not detected viscid fluid in the non-waxy pitcher types 4, 5, 6 and 11). Moran et al. (2012) also record visco-elastic fluid occurring in some waxy species, such as *N. tobaica* Danser, yet our checks of several accessions of this species showed it to be non-visco-elastic (Cheek pers. obs 2019). Gaume et al. (2016) record viscid fluid in *N.hemsleyana* Macfarl. (which we confirmed) which is also waxy, and we confirmed this from observations of live plants cultivated at Kew.

Species with type 2, non-waxy, narrowly funnel-shaped upper pitchers, can have type 1 waxy, ovoid-cylindric lower pitchers on the same plant e.g. *N.rafflesiana.* A key element of this stage-dependent heteromorphy (difference in shape dependent on stage of development) explains the loss of the waxy zone. This seems to be the switch in pitcher shape. Lower pitchers are usually ovoid-cylindric (also with the tendril uncoiled, placed in front of the mouth, and with the front of the pitcher facing the stem and having a pair of fringed wings). Upper pitchers instead, are more or less funnel-shaped in type 2 (and generally also face away from the stem, and have the tendril coiled and arising at the rear of the pitcher, and the fringed wings reduced to a pair of low ridges). This change in shape seems to be important in deciding whether or not the pitcher is waxy on the inner surface, or not. While species with narrowly cylindrical or ovoid-cylindric upper pitchers retain the waxy zone, those species which have funnel-shaped pitchers (widening gradually from base to apex) lack a waxy zone, or have only a vestigial, minute waxy triangle as described above. It is as though in the funnel-shaped pitchers the development of the upper cylindrical waxy zone is “switched off”, the ovoid basal non-waxy detentive zone instead expanding to fill the gap, and directly support the peristome instead. Gaume et al. (2016) report that in *N. rafflesiana* not only does the shape and waxiness of the pitcher switch from lower to upper pitcher, but that lower pitchers have non-viscid fluid, while upper pitchers have viscid fluid. In addition, the upper pitchers (type 2) of this species capture more flying insects than do their type 1 lower pitchers. In a study of six different *Nepenthes* species co-occurring at two sites in lowland Brunei, Gaume et al. (2016) found that pitcher shape critically influences the capture of flying insects, stating that “flying insects are clearly associated with funnel-shaped pitchers”.

In contrast species with waxy, narrow, cylindrical pitchers (e.g type 1 and type 10) proved more effective in trapping ants and termites respectively.

*Nepenthes madagascariensis* Poir. is exceptional in that it is the only species in the basal grade of the genus that does not have type 1 pitchers. Its narrow funnel upper pitchers, typical of those in type 2, were found to trap significantly more Coleoptera, Diptera and Lepidoptera (flying insects) “than in all other *Nepenthes* species” (Rembold et al. 2010). However, unusually for type 2, it does not have visco-elastic but watery fluid (Cheek pers. obs.). Bonhomme et al. (2011) hypothesised that the visco-elastic fluids in *Nepenthes, Drosera*, and *Drosophyllum* (in the last two genera appearing as insect-trapping mucilage on the leaf tentacles) have a common and thus a plesiomorphic origin. However, evidently the visco-elastic liquid trait lay dormant or was not re-acquired until after the genus arrived in Borneo. The earliest branching species of *Nepenthes* in which it occurs is *N. rafflesiana.*

Visco-elastic fluid is effective at retaining small prey, such as small ants e.g. *Oecophylla smaragdina* which are totally unable to free their bodies from it. However, larger ants, such as *Polyrhachis* species, can haul themselves out, and climb up the non-waxy pitcher wall in *Nepenthes rafflesiana* (Gaume & Forterre 2007).

However, species with waxy zones are far more effective at retaining prey at the bottom of their walls than those species with non-waxy walls. Comparing the similar and closely related *Nepenthes rafflesiana* var. *elongata* (that is *N. hemsleyana*) which has waxy walls, with *N. rafflesiana* var. *typica* (that is *N. rafflesiana*) which is non-waxy, Gaume & di Gusto (2009) found in experiment that the former retains 73.5% of trapped individuals, while the latter only 28.5%.

Bonhomme et al. (2011) quantified visco-elasticity and wax-levels of 23 cultivated species of *Nepenthes*, measuring their effects on the retention rates of flies and ants placed in the lower pitchers of 12 of these species. In the 23 species they found that none of the species with high levels of waxiness was found to exhibit a very visco-elastic fluid, and vice versa. They concluded that there were two strategies: a ‘waxy’ strategy and a ‘visco-elastic’ strategy. While retention rates for ants increased with waxiness and visco-elasticity, for flies retention rates did not depend on the amount of wax but were far higher when the fluid was visco-elastic. Two thirds of the 23 species in their study were classified as visco-elastic and these species tended to be montane rather than lowland. This agrees with the observations of Collins (1980) that in montane habitats in Borneo, ants are relatively fewer in number, but flying insects relatively more abundant.

**Pitcher type 3 (Fig. 3).**
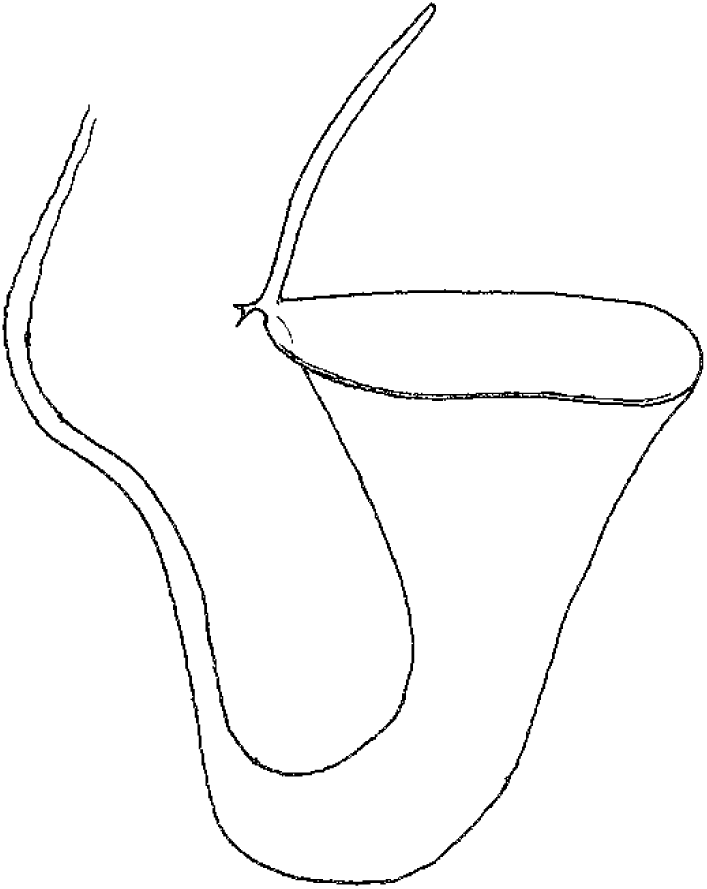
Flypaper (N. inermis Danser). This pitcher type, as with type 2 pitchers, have viscid fluid and lack an inner waxy zone, but in contrast, they are more diminutive, usually only 3-5 cm long, and much broader in comparison to their length, with a length: breadth ratio in the region of 1 or 1.5: 1. The upper part of the pitcher is usually a very broad, open shallow bowl that drains into the slender, cylindrical, fluid-filled lower part. In those species where observations have been recorded, small dipterans, such as midges and mosquitoes, are trapped in the bowl, as though by flypaper on the inner sticky, usually yellow-green inner wall of the pitcher which is moistened by the viscid pitcher fluid, the prey slowly sliding down into the column of fluid at the base of the pitcher, below the broad, funnel-shaped upper part. Previously we had considered that Wistuba (1994, related in Cheek & Jebb 2001: 82) was first to record how in the exemplar species *N. inermis*, rainwater entering the pitcher does not mix with the denser, insect-containing viscid fluid at the base, and is at intervals tipped out from the pitcher by the pitcher overbalancing due to the added weight. However, these very observations for this species were in fact first made by Kato et al. 1993. Kato et al. studied ten species in Sumatra, including *N. inermis* (mistakenly identified as *N. bongso* Korth.). Kato et al. stated that this species was particularly unique among the 10 studied, trapping few ants and many small midges. In fact, the majority, 60% of the prey were diptera (flies), some of which were adults of potential pitcher inhabitants. The fluid contained no living inhabitants. These facts suggested to Kato et al. that the pitcher may attract adults of phytotelmata inhabitants and trap them.

This prey-type and trapping mechanism has been recorded in the unrelated *N. eymae* Danser of the Regiae section in Sulawesi (Cheek & Jebb 2001: 82, McPherson 2009: 993). Two atypical species of Sect. Tentaculatae Cheek & Jebb in Sulawesi also have type 3 pitchers e.g. *N.pitopangii* Chi.C.Lee et al., and *N. undulatifolia* Nerz et al, and in New Guinea, N. paniculata Danser. Most of the type 3 pitcher species however occur with *N. inermis* in a group of Sumatran Montanae which were characterised in Cheek et al. (2017) as subsect. Poculae-Ovis (“the egg-cups”, due to their size and shape), so that type 3 pitchers have arisen independently in four different parts of the *Nepenthes* evolutionary tree.

**Pitcher type 4 (Fig. 4).**
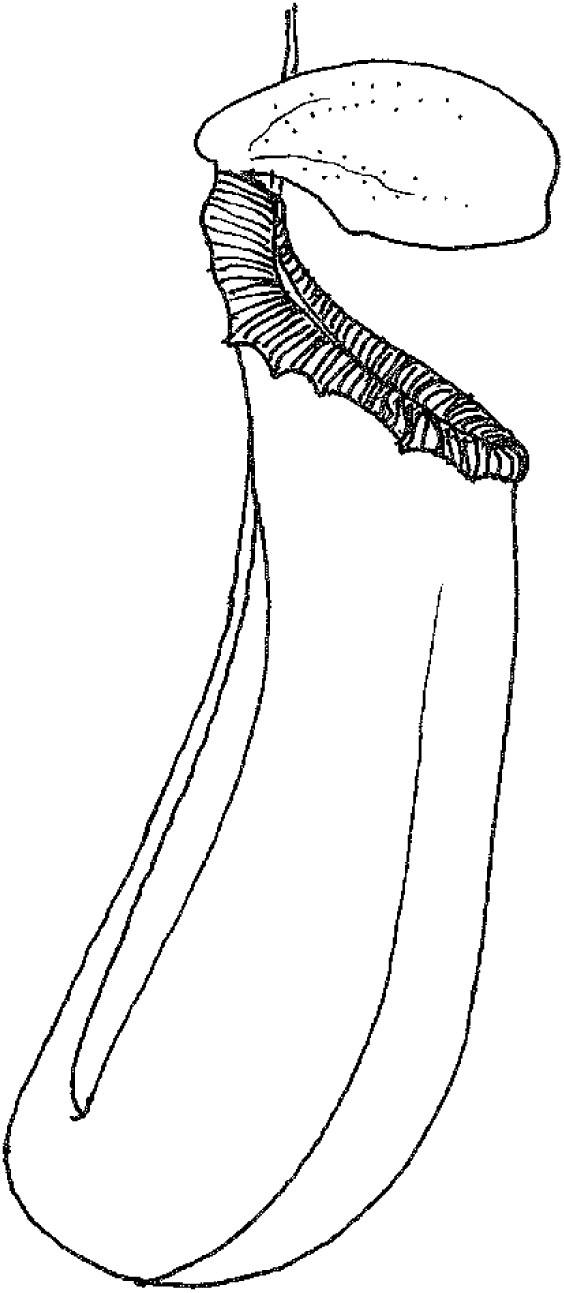
Stout cylinder. (*N. insignis*). These species bear stout cylindrical upper pitchers which lack a waxy zone, but which also lack visco-elastic fluid. They have a length: breadth ratio of c. 3: 1. They are most similar to type 2 pitchers, but lack the narrow-funnel shape of those species. Unlike type 6 pitchers, the peristome does not develop a distinct, vertical column. The species mostly have wide, deeply ridged peristomes e.g. *Nepenthes veitchii* Hook. f., *N. hurrelliana* Cheek & A.L. Lamb and *N. truncata* Macfarl., however *N. hirsuta* Hook.f. has a slender peristome. These four species are spread over three branches in the Nepenthes evolutionary tree (Murphy et a. 2019). The largest cluster of species occurs in a fourth branch: practically all of Sect. Insignes Danser, including *N. ventricosa* Blanco are type 4, and *Nepenthes bicalcarata* Hook. f. are placed here, The last is unique in its commensal relationship with the ant *Camptonopus schmitzii*. Prey composition and analysis of the trapping syndrome is available only for the last species. Gaume et al. (2016) found that *Nepenthes bicalcarata* is a generalist at their two Brunei study sites, each site with the same six species. Upper pitchersof *N. bicalcarata* predominantly trapped termites, while the lower pitchers predominantly trapped ants.

**Pitcher type 5 (Fig. 5).**
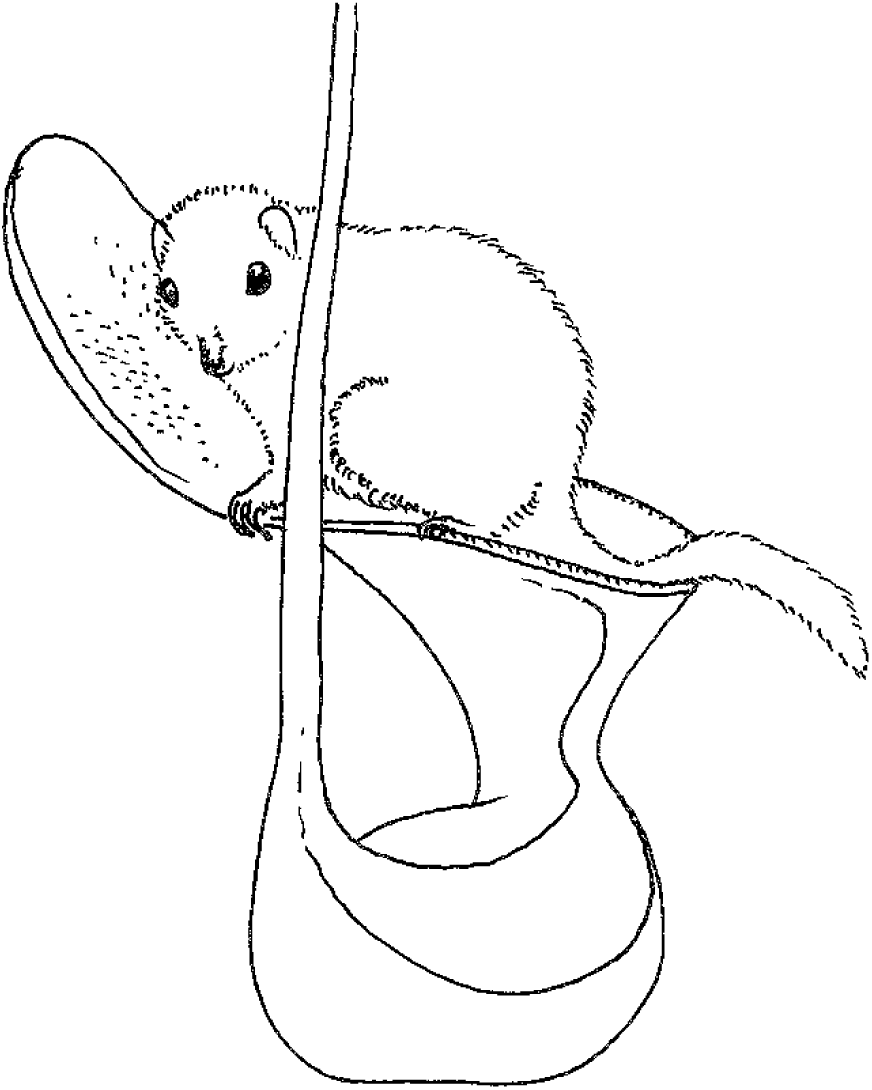
Tree shrew lavatory (*N. lowii* Hook.f.). Clarke et al (2009), Chin et al. (2010) and Greenwood et al. (2011) are among several studies documenting that the latest stage pitchers of *N. lowii N. rajah* Hook.f. and *N. macrophylla* (Marabini) Jebb & Cheek attract and trap the droppings of the tree shrew *Tupaia montana* and that these are a major source of nutrients for the species. It is speculated that the close relative of the first species, *N. ephippiata* Danser may also obtain nutrients in this way. Pitchers that are not full-sized, and large enough for the tree shrews to mount, feed and defaecate into depend instead on insect capture. While the upper pitchers of the first species appear not to trap animals at all, those of the second and third species are known to trap insects also. This pitcher type is characterised by a large, robust pitchers that can support the weight of the animals and which lack a waxy zone, but which also lack visco-elastic fluid. The mouth is large, c. 10 cm diam., with a peristome that facilitates gripping by all four limbs of the target animals that stand astride the pitcher mouth while licking nectar secreted by the lid angled at 90 degrees or more from the plane of the mouth. In *N. rajah*, a species of rat has also been recorded defaecating in the pitchers. The several records of drowned rats being found in pitchers of these species may relate to deaths while the animals were attempting to feed. In *N. lowii* (and possibly *N. ephippiata*) upper pitchers are visited by the animals, while in the *N. rajah* and *N. macrophylla*, upper pitchers are not, or rarely, formed. *Nepenthes attenboroughii* A.S.Rob. et al. of Palawan also has lower pitchers that meet the criteria of type 5, and although they trap arthropods, they are also visited by tree shrews, and have been recorded trapping the same (Mey 2013), although defaecation has not yet been recorded. Similarly, *Nepenthes merrilliana* Macfarl. of Mindanao also has pitchers that fit the specification of type 5, although tree shrews have not been recorded visiting and the lid angle in some traps does not exceed 90 degrees. Since pitchers with a mouth diam. of <c.10 cm cannot benefit from mammal nutrients, this factor seems likely to have contributed to selection for larger pitchers in the species concerned.

**Pitcher type 6. (Fig. 6).**
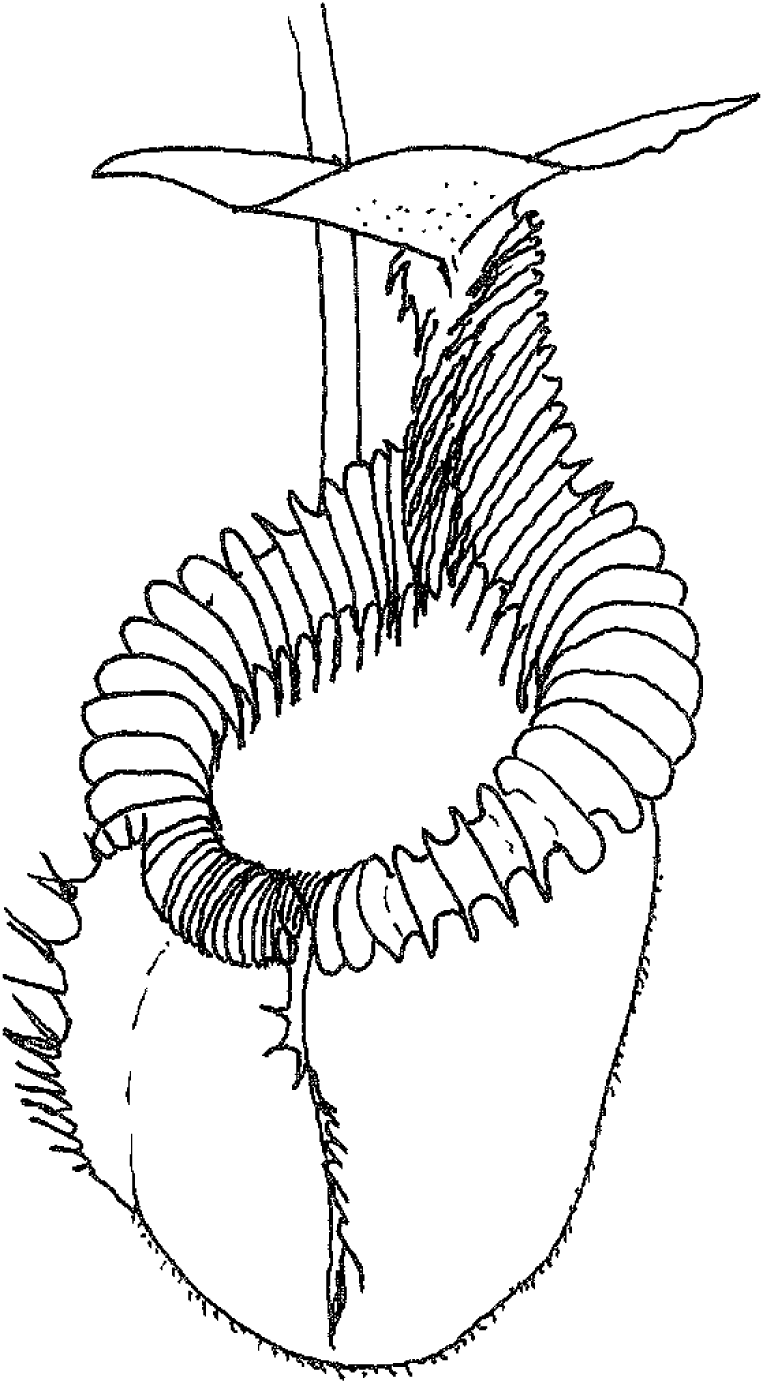
Globose-hairy (*N.mira* Jebb & Cheek). Globose-hairy or type 6 pitchers lack a waxy zone, lack visco-elastic fluid, and are usually large and shortly cylindrical to globose Upper pitchers are either not, or rarely formed. The lids lack appendages and are held on a well-developed and distinct vertical column formed from the peristome which extends to the lower lid surface. The peristome ridges can be developed into blade-like wings (e.g. *Nepenthes villosa* Hook.f., N. mira). The outer surface of the pitchers often have long, early caducous hairs.

This distinctive pitcher type was previously considered to represent a closely related, natural group of species. The term villosa group or complex was coined after the discovery of *N. mira* from Palawan when it was placed with *N. villosa*. of Kinabalu and its relatives in NE Borneo due to their morphological similarities. to these were added the minute but morphologically similar *N. argentii* Jebb & Cheek of Sibuyan (Cheek & Jebb 1999). The term villosa group or complex was adopted and expanded by Robinson et al. (2009) to include two newly discovered species from Palawan, and also *N.peltata* Sh.Kurata of Mindanao. However, the molecular phylogenetic work of Murphy et al. (2019) showed that the NE Borneo, Palawan, Sibuyan and Mindanao species referred to above fall in four separate clades so that their morphological similarity must be due to convergence. However, the source of nutrition targeted by species with this type of pitcher is not known. All species are restricted to high altitude ultramafic habitats.

**Pitcher type 7. (Fig. 7).**
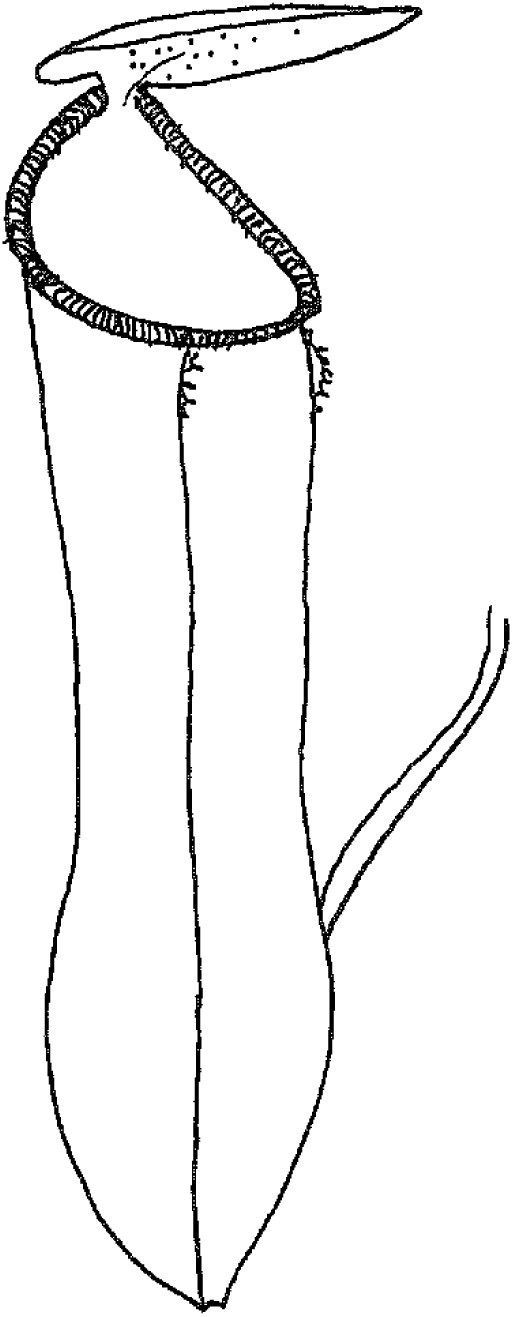
Flick of the lid (*N. gracilis* Korth.). Bauer et al. (2012b) reported how, in this species, ants are trapped by a mechanism previously unreported in *Nepenthes.* The lower surface of the lid, uniquely in the genus so far as is known, has waxy platelets as in the inner surface of the upper pitcher of type 1 species. Ants, feeding on the nectar produced from the large, but few nectar glands placed on the lower lid, have no difficulty maintaining their footing upside down. That is, until large tropical raindrops hit the upper surface of the lid. Such events, due to the elasticity of the lid hinge, “flick” the lid downwards sharply, and the ant, its footing security impeded by the wax platelets, is propelled down towards the detentive zone. *N. gracilis* otherwise would be regarded as a type 1 species since it has waxy, cylindrical or ovoid-cylindrical pitchers. However, some other species are postulated to have the same mechanism although they lack wax on the lower surface of the lid. In these species the presence of hairs on the surface, rather than of wax, is hypothesised to reduce the security of footing of insects such as ants. Those species with such hairs are *N. maryae* Jebb & Cheek from Sulawesi, of the Tentaculatae group, *N. oblanceolata* Ridl. (Regiae group, New Guinea) and *N. macfarlanei* Hemsl. (Montanae, Peninsular Malaysia), (Cheek & Jebb 2016). All these species, but for their lower surface lid modifications would otherwise be included in type 1, having waxy, cylindrical pitchers with watery fluid. Gaume et al. (2016) found that c.98% of the prey of *N. gracilis* consists of ants, each pitcher trapping over 100 individuals, a higher percentage than for any of the other five species growing at the same study sites in Brunei.

**Pitcher type 8 (Fig. 8).**
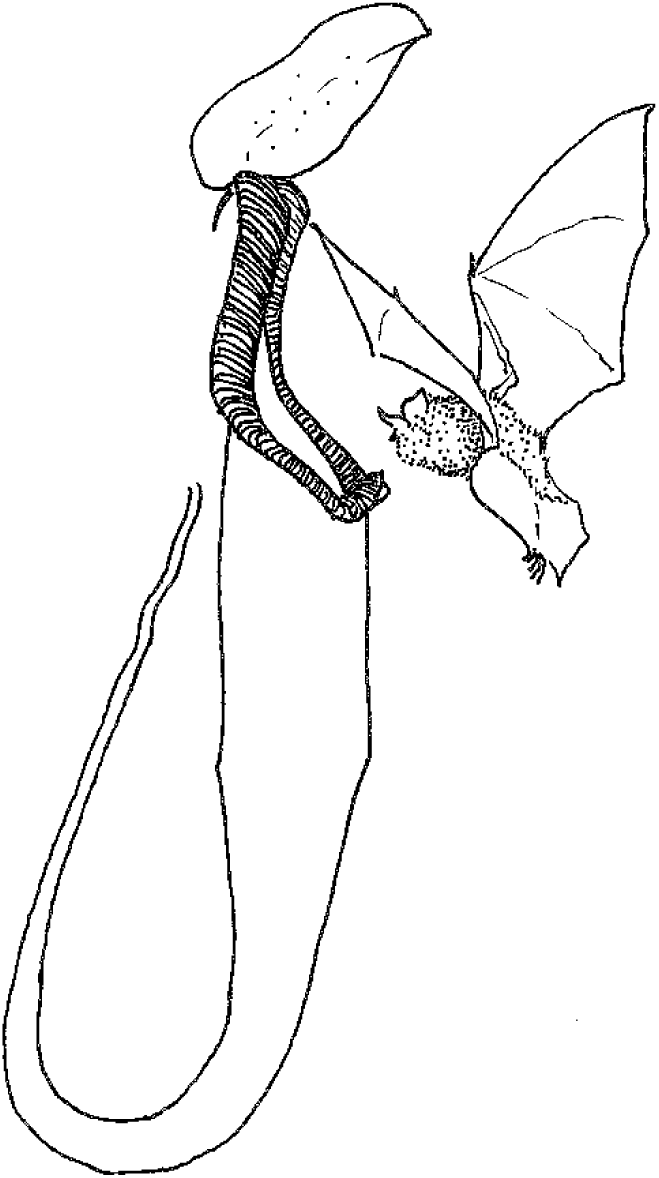
Bat-roost (*N. hemsleyana*). Grafe et al. (2011) reported how this species secures nutrients predominantly from the droppings of *Kerivoula hardwickii*, Hardwickes’s bat, which roosts within its pitchers. *N. hemsleyana* was confused with and synonymised by Danser (1928) with *N.rafflesiana.* Several papers have reported how *N. hemsleyana* has lost the characters of scent, and nectar production seen in other insect-trapping species (e.g. Gaume et al. 2016). It has developed a feature at the mouth of the pitcher that acts as a parabolic beacon, enabling bats to locate their potential roosting sites with ease by echo-location (Schöner et al. 2015). Although it gains nutrients mainly from bat droppings, *N. hemsleyana* still retains the ability to trap insects. Gaume et al. (2016) reported that upper pitchers contained about 25 individual prey items, mainly ants. In comparison, pitchers of *N. rafflesiana* at the same site and time contained about 100 prey items. The bats sometimes also roost in pitchers of *N.bicalcarata*, which are more frequent, but which are less preferred. Bats roosting in *N. hemsleyana* were found to have higher body weight and fewer parasites than those in *N. bicalcarata* (Schöner et al. 2013). Apart from hosting bats, morphologically *N. hemsleyana* has type 1 pitchers, standing apart from other such species only by its highly elongated pitchers. These are similar in dimensions to those of *N. spectabilis* Danser of Sumatra which can be postulated also to be used as bat roosts. This should be tested with field observations. Might other species of *Nepenthes* also have such mutualistic relations with bats? Since the bat mutualism, and even the identity, of *N. hemsleyana* was overlooked, despite intensive study of animal-*Nepenthes* interactions in its range in Brunei by many scientists over decades. Hardwicke’s bat occurs from India to Indonesia

**Pitcher type 9. (Fig. 9).**
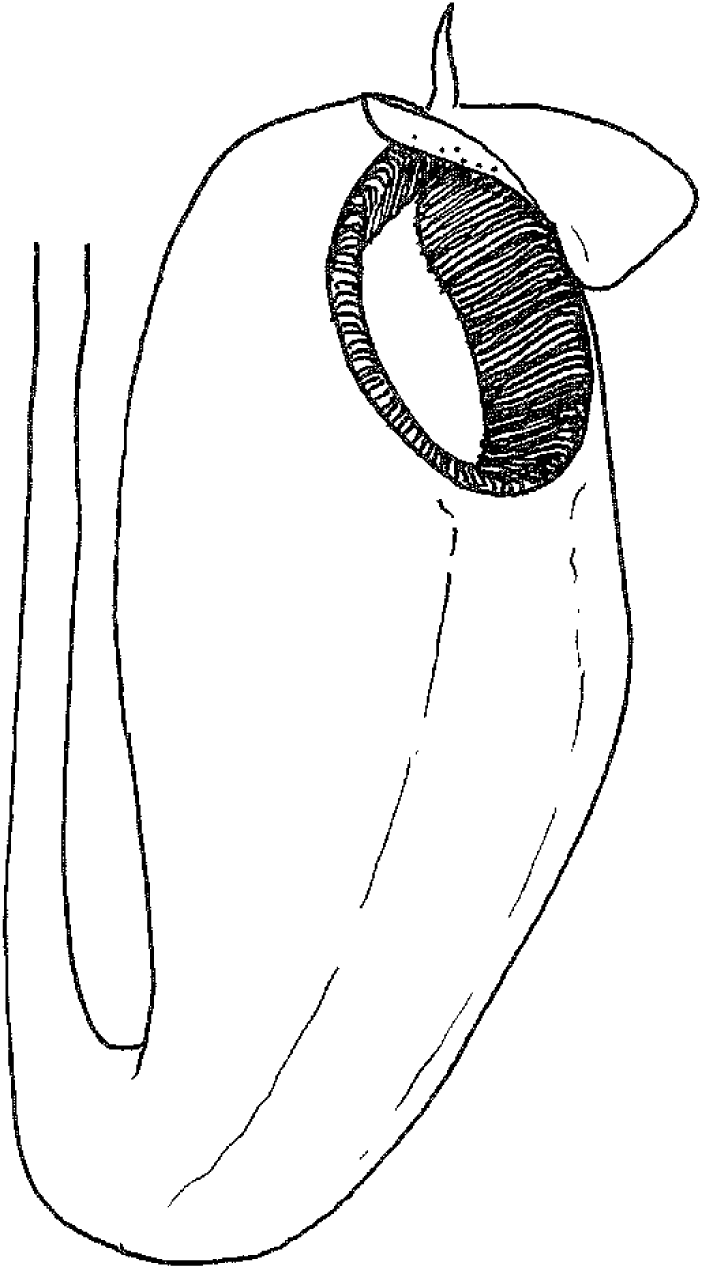
Light-trap (*N.aristolochioides* Jebb & Cheek). Pitchers with the mouth at the apex, held more or less horizontally and facing the sky are usual in the genus and are present in all other pitcher types. However, in two species, *Nepenthes klossii* Ridl. (New Guinea, Sect. Regiae) and *N.aristolochioides* (Sumatra, Sect. Montanae subsect. Poculae-ovis) the pitcher resembles a bladder (Jebb & Cheek 1997) and the mouth faces horizontally while the top of the pitcher is a windowed dome, the light panels lacking pigment, creating a bright translucent window within the dark peristome. A similar construction is present in the American pitcher plants of the Sarraceniaceae: *Darlingtonia californica* Torr., *Sarracenia minor* Walt. (as pointed out by Jebb 1991), and *Sarracenia psittacina* Michx. (Schaefer & Ruxton 2014). In laboratory conditions Moran et al. (2012) showed with *N. aristolochioides* that flies of *Drosophila melanogaster* arriving under the opaque lid, which shades the mouth, land on the vertically held peristome, feed on the nectar, and depart towards the light window at the rear and top of the pitcher, becoming stuck to the non-waxy wall and due to the viscid fluid slowly slide down towards the base where digestion occurs. If the rear of the pitcher is shaded, lower numbers of flies are trapped than otherwise, confirming that the flies enter the pitcher and are then trapped due to attraction by the lighted windows. This species is very closely related to seven others in its subsection which are placed in type 3. The identical structure is seen in *N. klossii*, except that the inner walls are waxy, and not sticky, and the fluid is not known to be viscous. In both species, flies are assumed to be the main prey items, but this needs verification.

**Pitcher type 10. (fig. 10).**
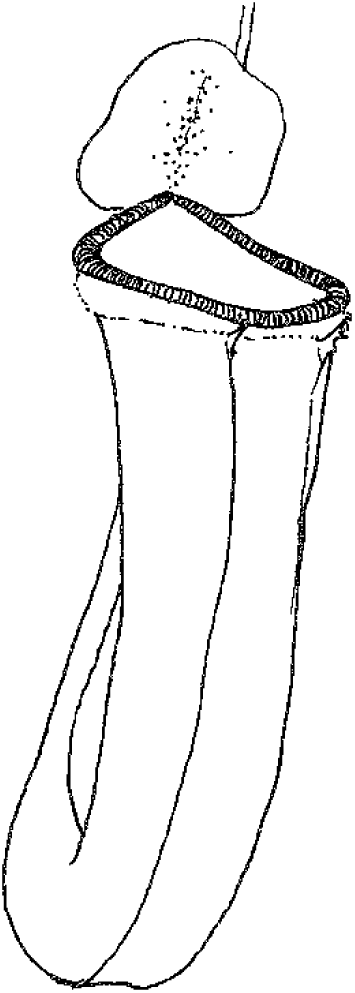
Termite trap (*N. albomarginata*). Only this single species is generally known to specialise in trapping termites, reported from its range in Borneo (Merbach et al., 2002), Sumatra and Peninsular Malaysia in numerous publications. Kato et al. (1993) appear to be the first to have reported this phenomenon in *N. albomarginata* in a study of ten *Nepenthes* species from Sumatra. In that study, 60 % of the prey recorded in that species were termites. Similarly, Adam (1997) in Borneo reported that 86% of trapped were termites. While it is widely reported that the crucial character for termite trapping in *N. albomarginata* is the edible hairs in the band on the outer surface of the pitcher below the peristome, Gaume et al. (2016) in a study of six species in lowland Brunei report that additional characters that favour trapping of termites are cylindrical, narrow diameter pitchers. This was because in their study in lowland Brunei, high incidence of termite trapping was also recorded in the upper (but not the lower) pitchers of *Nepenthes bicalcarata* (not included here in type 2) which has these features, but which lacks the band of white edible hairs of *N. albomarginata.*

**Pitcher type 11. (Fig. 11).**
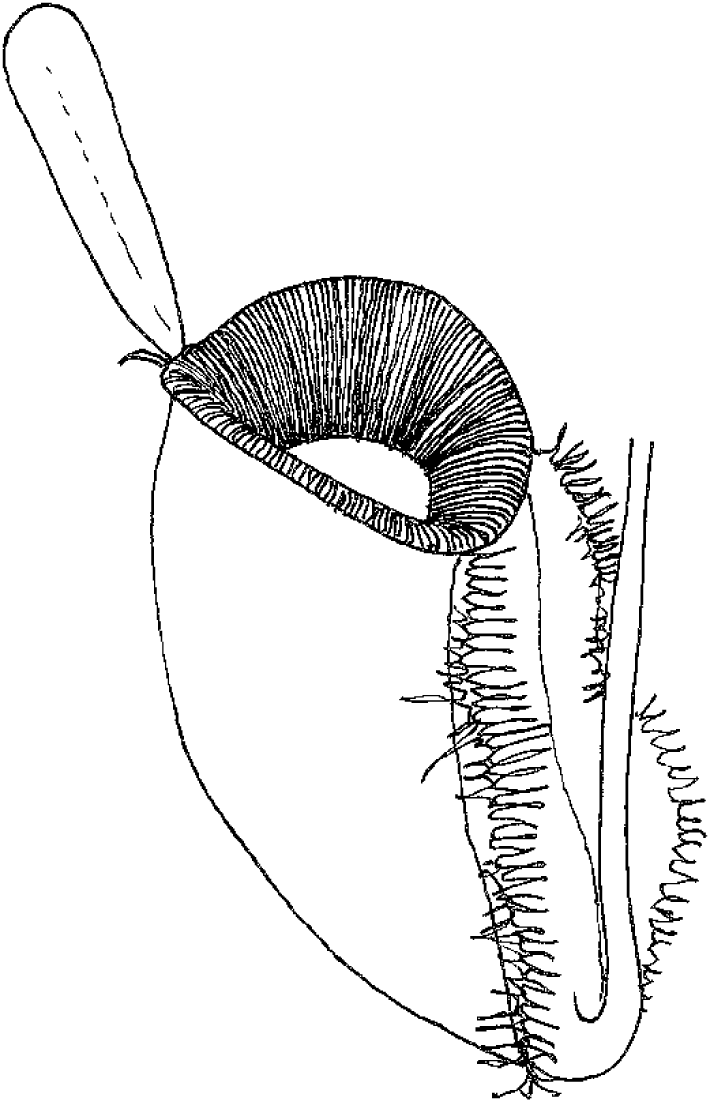
Pitfall (*N. ampullaria* Jack). At present this is the single species which, although trapping low numbers of ants (about 10-15 per plant (Gaume et al. 2016)), predominantly traps and depends nutritionally upon leaf-litter e.g. Cresswell (1998), Pavlovič (2011). The lack of insect-attractive colouration (pitchers are predominantly green), scent and nectar (important in attracting insects such as ants) are seen to reduce its efficiency at trapping insects. The prolifically produced, wide-mouthed but squat pitchers, produced in carpets on the forest floor, have lids which are reflexed, facilitating trapping of litter material falling in from the forest canopy above. No other species is known to have these attributes. In other respects, Pitcher type 11 is similar to types 4-6 in lacking both a waxy surface and visco-elastic fluid.

**Pitcher type 12. (Fig. 12).**
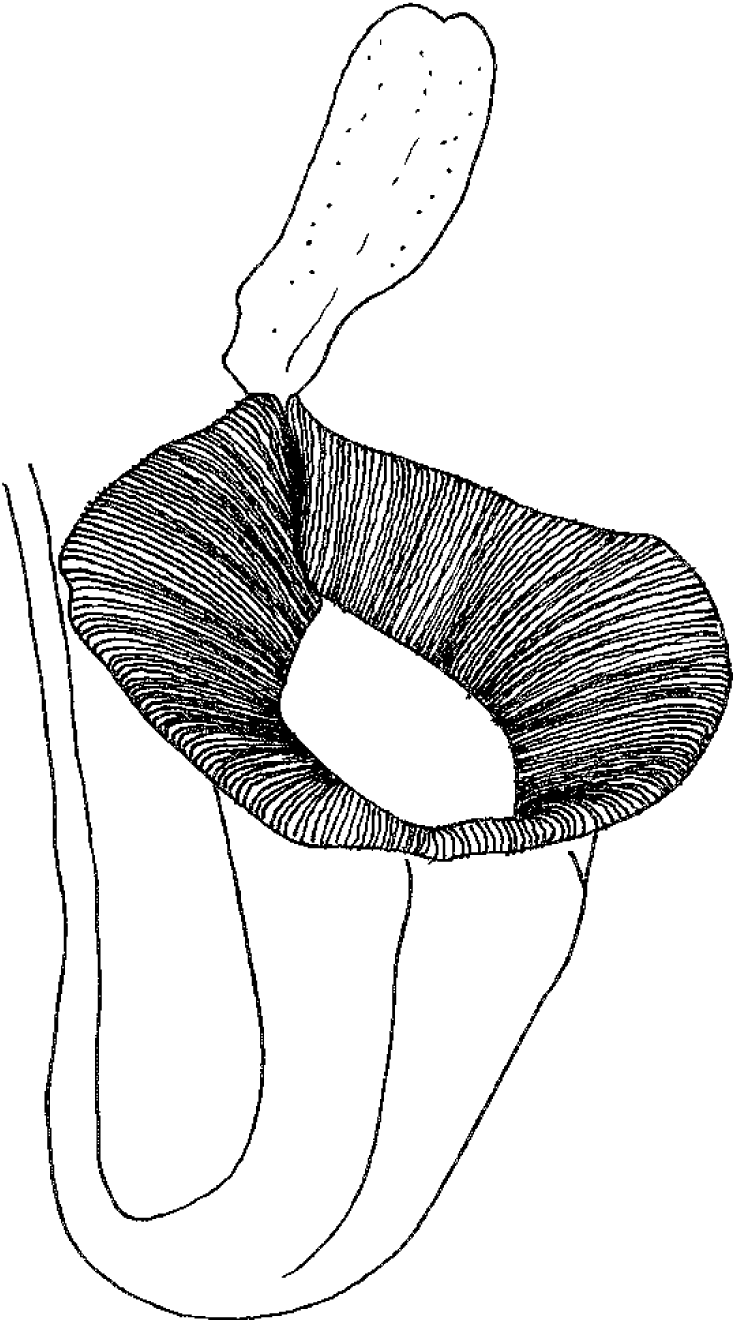
Flat lip (*N. jacquelineae Nepenthes jacquelineae* (Sumatra, Sect. Montanae) and *N. platychila* Ch.C.Lee (Borneo, Sect C.Clarke et al.).. Regiae, Lee 2002) both have all the features of type 3 pitchers: widely funnel-shaped to cup-shaped, non-waxy upper parts of pitchers draining into a more slender cylindrical lower part with viscous liquid. Both species however differ from all type 3 species in their very large, flat, dark red peristomes which can be up to 3.5 cm wide in *N. jacquelineae.* These peristomes have been speculated to act as a landing platform for large flying insects such as blattid cockroaches and moths, which might act with the contrasting lighter, green pitcher body as a light trap for such prey. It is also possible that such prey are lured into a precarious place above the mouth by the copious nectar produced from the lower surface of the lid from large nectar glands 1.5 mm diam. (Clarke 2001). The morphological convergence between the pitchers of these two species is remarkable. Further investigation of the prey trapped in the wild would be desirable for both species. *Nepenthes echinostoma* Hook.f. is a species closely related to *N. mirabilis* but with a remarkably flat and extended peristome which resembles type 3, although it has a different pitcher shape and inner surface

## Discussion

Further pitcher types are likely to be discovered in *Nepenthes* as more species are discovered, and as those that are known are better researched. For example, *Nepenthes bicalcarata* here included in pitcher type 4, differs from all other species ascribed to that pitcher type (and from all other *Nepenthes*) in a) its mutualism with the specialised ant *Camptonopus schmitzii* which it accomodates, and b) in its fang-like peristome extensions, so that it may prove to represent a further trapping syndrome.

There are few studies that compare trapping of prey by different species of Nepenthes with different pitcher types at a single site. Among the ten species in Sumatra studied by Kato et al. (1993) three species, equating to different pitcher types, were sympatric at in the submontane forests of Gunung Gadang. These were, *N. inermis* (type 3), *N. spathulata* Danser (type 2) and *Nepenthes* B (unidentified, probably type 1). All three had different prey assemblages. The availability of prey was thought to be largely similar among the three species because their microhabitats were largely similar. This suggested to Kato et al. that the differences between prey assemblages was due to different prey trapping patterns.

A second study by Gaume et al. (2016) in lowland heath forest of Brunei studied six species, each with a different pitcher type: *N. rafflesiana*, type 2; *N. bicalcarata*, type 4; *N. gracilis*, type 7; *N. hemsleyana*, type 8; *N. albomarginata, type 10 and N. ampullaria*, type 11). All six species were present at two sites. They excluded from consideration *N. hemsleyana* (bat droppings, type 8) and *N. ampullaria* (litter, type 11) since they are largely non-carnivorous. Their analysis stated that there were three main carnivorous syndromes A) The “flying insect syndrome”, characterized by funnel-shaped pitchers of large diameters, with a yellow dominant colour, an acidic viscoelastic fluid, nectar secretion, and the delivery of a sweet scent; B) The “ant syndrome”, which is less specific and is characterized primarily by nectar secretion, then by fluid acidity and, to a lesser extent, a waxy trap; and finally C) The “termite syndrome”, characterized by narrower pitchers and a shape that is closer to a cylinder, with non-viscous fluids. Termite capture is also greatly enhanced by the presence of a rim of edible trichomes or the symbiotic presence of the hunter ant, *C. schmitzi.* The flying insect syndrome clearly maps onto N. *rafflesiana* (type 2), while the termite syndrome maps onto type 10 (*N.albomarginata*), and to some extent also type 4 (*N. bicalcarata*). Ant trapping is more widely spread among the species, but linked especially with *N. gracilis*, although Gaume et al. appear unaware of the “Flick of the Lid” mechanism revealed by Bauer et al. (2012b). These two studies, in different geographic locations, in different habitats at different altitudes, indicate that sympatric species of *Nepenthes* have different prey assemblages, and different trapping mechanisms. Additional studies are needed to test whether this is always the case.

Additional cryptic pitcher traits that are likely to be important in trapping syndromes, but for which both comparative and experimental data are absent or very sparse are: a) pitcher fragrance (Di Gusto et al. 2008); b) acidity levels of pitcher fluid (eg. Gaume et al. 2016); c) enzymatic differences in the pitcher fluid (Biteau et al. 2013); d) lower lid nectar gland and appendage structures (e.g. Cheek 2015); and e) peristome ridge and teeth morphology.

Pitcher morphology might not always be related solely to nutrient trapping. Water storage as a buffer for dry periods has not been adequately researched as a potential function of pitchers.

## ACKNOWLEDGEMENTS

Thanks to the past and present staff of the tropical nursery, especially Kath King, Nick Johnson, James Beattie, Rebecca Hilgenhof and Tom Pickering for maintaining the live Nepenthes collection that has been so important for our ongoing studies of *Nepenthes.* We thank Tim Barraclough, Magdalen College, University of Oxford for suggesting that we categorise *Nepenthes* species by pitcher type.

