## Supplemental Table 1. for "A classification of functional pitcher types in *Nepenthes* (Nepenthaceae)"

| **Species** | **Visco-elastic liquid?** | **Wax?** | **Assigned Pitcher Type** | **Accession no.** |
| --- | --- | --- | --- | --- |
| N. rajah |  |  | 5 | 1982-144 |
| N. ephippiata |  |  | 5 | 2004-2400 |
| N. lowii |  |  | 5 | 2004-2412 |
| N. ampullaria |  |  | 11 | 2005-2130 |
| N. hemsleyana | X | X | 8 | 2005-1767 |
| N. bicalcarata |  |  | 4 | 2002-3114 |
| N. burkei |  |  | 4 | 2001-1418 |
| N. ventricosa |  |  | 4 | 2001-1417 |
| N. albomarginata |  | X | 10 | 2002-892 |
| N. madagascariensis |  |  | 2 | 1989-681 |
| N. maxima | X |  | 2 | 2004-2389 |
| N. mirabilis |  | X | 1 | 1981-5655 |
| N. tobaica |  | X | 1 | 2004-2386 |
| N. merrilliana |  |  | 5 | 2005-2114 |
| N. reinwardtiana |  | X | 1 | 2003-2777 |
| N. truncata |  |  | 4 | 2005-2188 |
| N. veitchii |  |  | 4 | 1990-928 |
| N. boschiana |  | X | 1 | 2009-826 |
| N. eymae | X |  | 3 | 1983-4248 |
| N. veitchii |  |  | 4 | 2005-1759 |
| N. fusca | X |  | 2 | 2003-2750 |
| N. rafflesiana | X |  | 2 | 2001-1391 |
| N. zygon |  | X | 1 | 2004-2413 |
| N. spectabilis |  | X | 8 | 2004-2387 |
| N. sibuyanensis |  |  | 4 | 2001-1848 |
| N. sanguinea |  | X | 1 | 2005-2181 |
| N. petiolata |  | X | 1 | 2009-264 |
| N. copelandii | X |  | 2 | 2005-1754 |
| N. stenophylla |  |  | 2 | 2004-2388 |
| N. gymnamphora |  | X | 1 | 2004-2411 |
| N. aristolochioides | X |  | 9 | Glasnevin s.n. |

Table 1. Species of Nepenthes scored for presence (X) (versus absence) of visco-elastic pitcher fluid, wax on inner pitcher surface (versus absence), pitcher type (1-12 see Results), and RBG, Kew living collection accession number. All material cultivated at the Royal Botanical Gardens, Kew (excepting N. aristolochioides from National Botanic Garden, Glasnevin, Ireland), all sampling from latest stage pitchers.
